# Inflammation as a Silent Partner in Opioid Addiction: A Regulatory Logic Model

**DOI:** 10.1101/2022.04.26.489572

**Authors:** J. Tory Toole, Matthew Morris, Cole A. Lyman, Garry Spink, Gordon Broderick

## Abstract

The brain and body consist of complex networks of interconnected feedback and feed forward loops. Because these networks are capable of supporting multiple homeostatic states, a stressor or combination of stressors may cause the network to become “stuck” in a persistent maladaptive state, for example, chronic pain and the potentiation of opioid dependency. The current research uses automated text mining of over 14,000 publications to assemble a regulatory circuit consisting of 44 immune and neurotransmission mediators linked by 188 documented regulatory interactions. Decisional logic parameters dictating the regulatory dynamics available to each network model were estimated such that predicted behavior would adhere to observed pathologies. Analysis of this psycho-neuroimmune network confirmed that a broad family of behavioral kinetics may be equally capable of supporting dynamically stable conditions of chronic pain, persistent depression and addiction behaviors. Despite differences in the predicted course of onset, these models typically point to characteristic patterns of increased inflammatory activity in the brain for each of these pathologies, specifically increased expression of the protein complex NF-kB and inflammatory signaling proteins IL1-B, IL6, and TNF. Potential treatments targeting both addiction and chronic pain may therefore benefit from the use of anti-inflammatory drugs as pharmacological potentiators of current behavioral interventions.

**Clinical Relevance:** This work establishes a methodology for understanding both illness-specific and shared mechanisms underlying addiction, chronic pain, and depression, and the corresponding expression profiles of psychoneuroimmune markers that might facilitate screening and treatment design.

## I. Introduction

Since 1999, around half a million people in the U.S. have died from opioid overdose, with each year seeing more deaths than the last. According to the CDC, this opioid crisis has occurred in three distinct waves, beginning with the increased therapeutic use of opioids in the 1990s. Since 1999, there has been a relatively steady increase in overdose deaths involving prescription opioids (natural and semi-synthetic opioids and methadone). The year 2010 saw the beginning of the second wave, which included an exponential growth in overdose deaths involving heroin. Finally, the third wave began in 2013 with an even sharper increase in overdose deaths involving synthetic opioids, particularly those involving illicitly-manufactured fentanyl (IMF) [1][2][3]. Despite the risks of addiction and overdose, many still prescribe opioids as a means of alleviating pain and suffering for those individuals seeking treatment. This, in a large part, is due to the overwhelming complexity of chronic pain. It is estimated that more than 30% of Americans have some form of acute or chronic pain [4][5]. U.S. retail pharmacies dispensed 245 million prescriptions for opioid pain relievers in 2014 [6][7]. For acute pain, opioids can be very effective and improve function. However, their benefits for relieving chronic pain are more questionable [8]. The etiology and mechanisms underlying both chronic pain and addiction are numerous and can be very difficult to pin down. They are best understood from a broad biopsychosocial perspective that effectively integrates neurophysiological factors as well [9]. The complex networks of interconnected feedback and feed-forward loops in the physiology of the brain and body give rise to various daily functions and behaviors. Some of these functions, such as increased anxiety, depressive behaviors, or fight-or-flight responses serve to help cope with stress or trauma. Because these neurophysiological networks are capable of supporting multiple homeostatic states by virtue of their complexity, a stressor or combination of stressors can, under the right conditions, cause the network to become “stuck” in a persistent maladaptive state, for example, chronic pain or opioid dependency.

In this paper we analyze a model psychoneuroimmune regulatory network consisting of 44 components, including neurotransmitters, immune mediators and behavioral constructs, to determine this network’s natural ability to stabilize in a maladaptive pattern, namely one potentiating chronic pain, addiction, and depression. We then are able to predict patterns of shared physiological markers that might be expressed across and between these three states. Such predictions have the potential to inform on possible molecular panels for the identification of molecular and behavioral risk factors as well as potential solutions for destabilizing these chronic conditions and promoting a return to a healthy homeostatic regulation in both chronic pain and opioid addiction.

## II. Methods

### A. Network Model Creation

In this work Pathway Studio^*^ (Elsevier Amsterdam) was used as a visualization tool to exhaustively search Elsevier’s s Knowledge Graph, a database of causal and associative interactions mined from over 8M full text journal articles using the natural language processing (NLP) engine MedScan [10]. The current research uses the text mining of over 14,000 peer-reviewed publications to assemble a regulatory circuit consisting of 44 immune and neurotransmission mediators linked by 188 documented interactions.

Once the network was created, a manual model check was performed to confirm the directionality of regulation effects in regards to the psychological and behavioral variables. The model included physiological markers implicated in chronic pain and opioid addiction, as well as markers from previous work on the underlying mechanisms of depression and anxiety [11]. The final version of the model can be seen in Fig. 1. Parameters dictating the decisional dynamics of the network were estimated such that predicted behavior would adhere to observed pathologies. These were described qualitatively based on domain expertise and stated as higher or lower than normal expression of a behavioral construct or physiological marker as discussed below.

**Figure 1:**
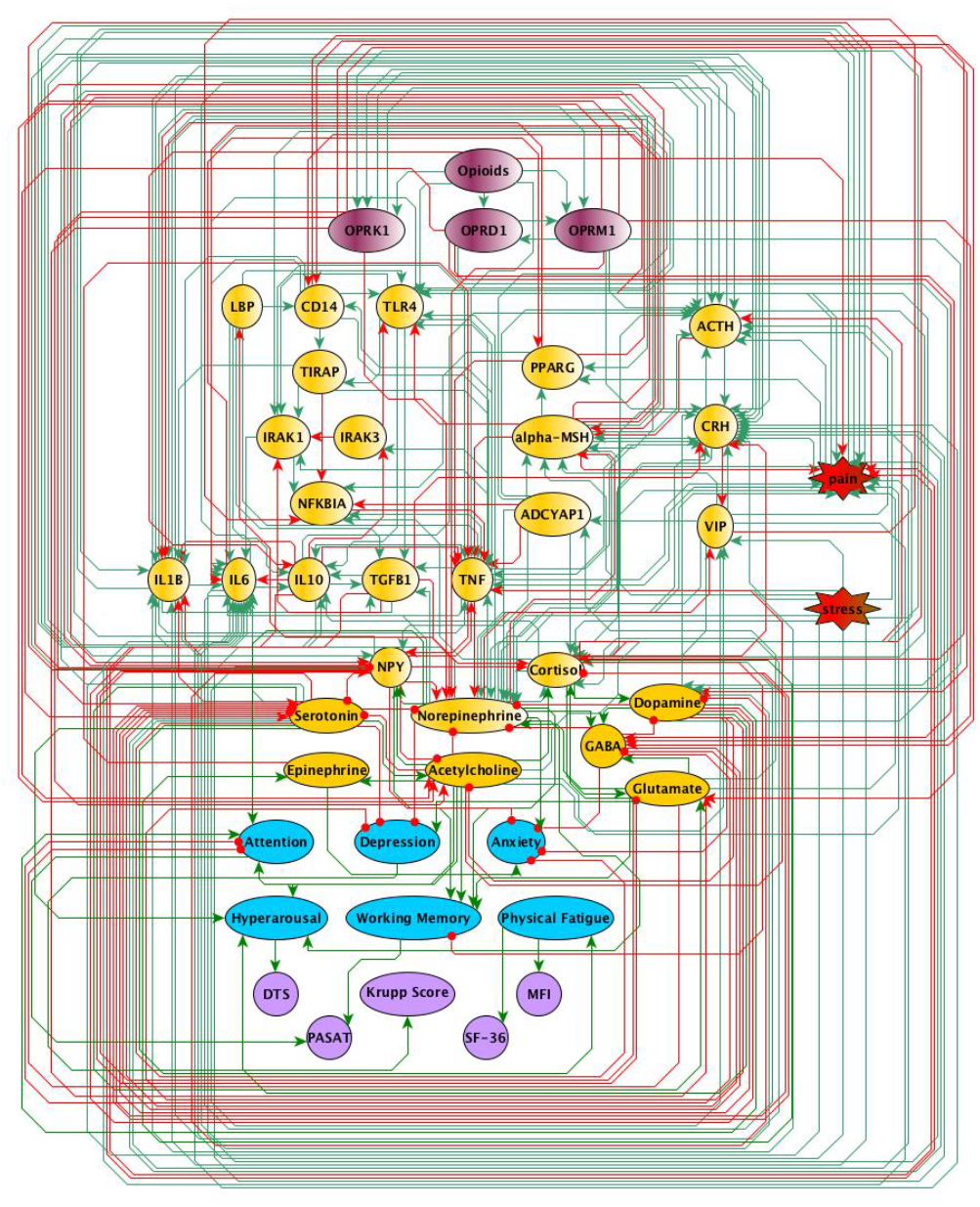
Neuroinflammation network model. Created through Pathway Studios, this model involves directional effects between physiological markers implicated in chronic pain and opioid addiction, as well as previous work on the mechanisms underlying depression and anxiety.

### B. Model Constraints

The set of feasible values for the logic parameters that dictate the model system’s dynamic response were selected such that they supported the prediction of resting states that would be consistent clinical observations in depression, chronic pain and addiction (Fig. 2**) [**12]. We hypothesized that the depressive state would, at a minimum, be constrained by high and low markers that were reported in previous work [11]. Additionally, we hypothesized that a “chronic pain” state would be minimally defined by “pain” being higher than baseline, and “opioids” not being administered into the system. A hypothesized “addiction” state would be minimally defined by chronically low dopamine, also with no opioids being administered into the system. Finally, an “opioid” state would be minimally defined as one in which “opioids” are chronically “on” and “pain” is persistently held in check at a “low” level. Finally, a “healthy” state was also hypothesized, as it should be the state to which the system is most adherent, and which would be characterized by all markers being expressed at their nominal levels “at rest,” and with the expression of behavioral markers such as depression and anxiety also being low. Of these five constraints, the hypothesized “opioid” persistent treatment state was not satisfiable, suggesting that the analgesic effects of opioids are not dynamically stable but rather are transient in nature and cannot be sustained indefinitely at least according to this regulatory model.

**Figure 2.**
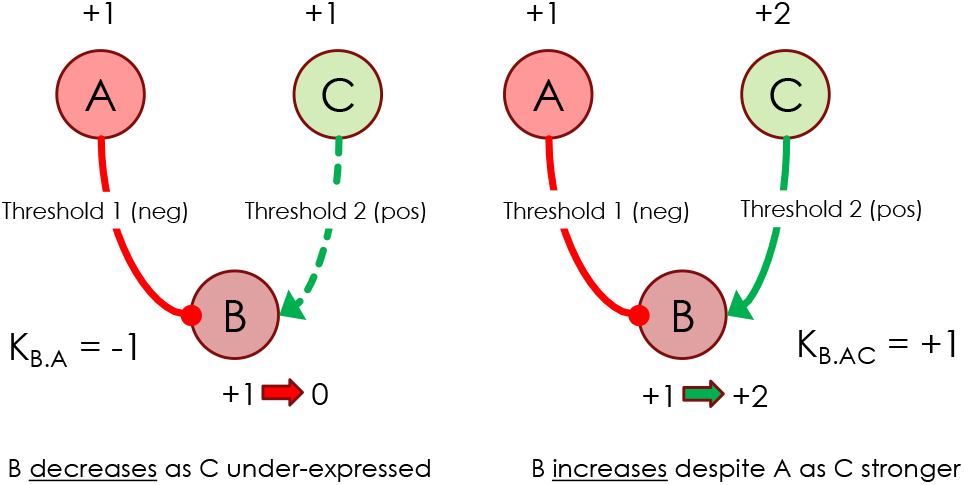
Discrete logic decisional parameters. In a basic circuit a node B can be downregulated by node A and upregulated by node C. In the left panel, when A is expressed at a state +1 in excess of its activation threshold whereas C is expressed at state +1 below its activation threshold, then node B is downregulated by node A acting alone (K_B,A_ = -1). In the right panel, when A is expressed at a state +1 in excess of its activation threshold but C is further expressed at state +2 above its activation threshold, then node B is regulated by both nodes. The net decisional weight of nodes A and C acting on B (K_B,AC_ =+1) is such that the net regulatory action is to upregulate B.

## III. Results

A total of 37,906 combinations of decisional logic parameters were found that support adherence of the circuit model to the minimally defined conditions described in section 2., However these collapsed into a much smaller set of 27 model classes each with similar dynamic behaviors. Despite producing slightly different response dynamics, all models unanimously predicted the same characteristic patterns of expression in the unconstrained and unspecified markers for each of the three stated pathologies. In other words all models agreed in their prediction of which markers were expressed at higher than normal levels, which were expressed at lower than normal, and which remained at nominal levels in each of the steady states. Interestingly, though some markers were characteristically expressed in a specific pathology, a surprising number were shared in at least 2 conditions. Of these, high acetylcholine and serotonin, combined with low expression of dopamine and GABA were common to both addiction and chronic pain. Similarly, predictions of high glutamate levels and low delta and mu opioid receptors were shared between addiction and depression. Finally, high levels of cortisol, IL1b, IL6, TGFB1, TLR4, and TNF, as well as low NFKBIA and PPARG were predicted to be shared by all three illness states. These results can be seen in Fig. 3.

**Figure 3:**
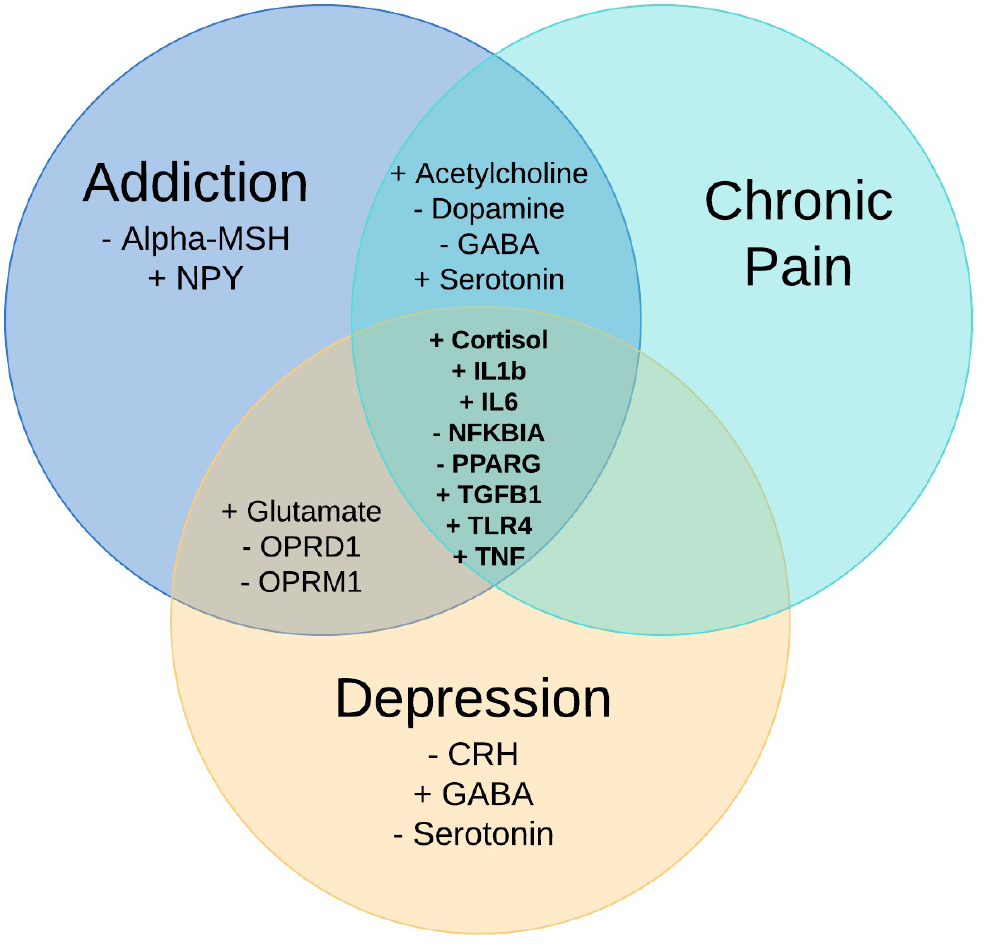
Persistent patterns of high and low marker expression. Overlapping expression patterns shared between the three illness states supported by the model. Shared markers include various neurotransmitters, hormones, and immune cells, and allow for some insight into possible network dynamics shared between these three illness states.

## IV. Discussion

In this paper, we attempt to capture the interaction dynamics of 44 neurophysiological markers and how they relate to depression, opioid addiction and chronic pain, illnesses that are extremely complex and difficult to treat. We utilized the Pathway Studios interface to search the Elsevier Knowledge Graph (Elsevier, Amsterdam), and effectively scanned the full text in tens of thousands of peer-reviewed articles, extracting putative cause-and-effect interactions between any two nodes of the network. This automated assembly of predictive models from prior knowledge is highly noteworthy as conventional manual curation of the literature remains by necessity much more restricted in coverage and ultimately suffers from significant observer bias. Integrating these regulatory circuits with a decisional logic that is constrained to reproduce experimental and clinical observations made it possible to simulate the behavior of the overall system and predict the detailed psycho-neuro-immunological profile of very cursory minimal definitions of depression, addiction and chronic pain. This model network correctly predicted the well-known gradual decrease in analgesic effects observed over the course of continued opioid use [13]. Most importantly, consensus among a broad family of competing models unanimously predicted an overlapping inflammatory signature shared between these pathologies. The connection between depression and immune system function has been documented for several decades [14][15]. Impaired immune function is a common and significant symptom of depression, and conversely, depression can sometimes be a symptom of various physical illnesses. More recent work even hypothesizes that depression can be thought of as a microglial disease intimately tied to immune function within the brain [16]. Furthermore, there is also recent evidence suggesting a link between immune dysfunction and addiction, including morphine treatment in rodents leading to increases in levels of proinflammatory cytokines such as IL-1b, TNF-a, and IL-6. Another study found that nuclear factor kappa light chain enhancer of activated B cells (NFkB), a transcriptional activator of inflammatory cytokines, can be overly activated by alcohol and other drugs of abuse [17][18]. The current work implicates specific immune targets that may be central to both addiction and depression, elucidating potential immune mechanisms that could not only further our understanding of these mental illnesses, but suggest treatment targets for future study. It is important to emphasize that this network of documented regulatory interactions combined with a very minimal qualitative description of these pathologies was sufficient to accurately predict the corresponding immune profiles despite that no actual experimental data was applied directly.

By harnessing prior knowledge in this way this particular methodology extends beyond statistical association and provides a glimpse into the possible mechanisms that underlie and drive illness. Indeed, understanding the overlaps between the addiction, chronic pain, and depression illness states may inform on the underlying regulatory processes governing their persistence, and support the design of possible strategies for medically assisted behavioral therapies. Ongoing work includes the use of large-scale Monte Carlo simulations to delve into the feasibility and likelihood of therapeutically redirecting this psycho-neuro-immune regulatory regime from one persistent resting state to the other, with the intention of determining how to most efficiently undermine illness in favor of a reliable return to health.

## Acknowledgment

This work was supported by Rochester Regional Health in conjunction with Elsevier BV (Amsterdam) under a collaborative research sponsorship (Broderick, PI). The authors would also like to thank Drs. Chris Cheadle, Hongbao Cao, Dicle Hasdemir, Anton Yuriev, Philipp Anokhin, Umesh Nandal and Thibault Geoui of Elsevier BV for their technical assistance and many helpful discussions.

## Mandatory Disclaimer

The opinions and assertions contained herein are the private views of the authors and are not to be construed as official or as reflecting the views of the Rochester Regional Health or Elsevier BV.

## Footnotes

* Copyright © 2020 Elsevier Limited except certain content provided by third parties.Pathway Studio is a trademark of Elsevier Limited.

